# TCRex: detection of enriched T cell epitope specificity in full T cell receptor sequence repertoires

**DOI:** 10.1101/373472

**Authors:** Sofie Gielis, Pieter Moris, Wout Bittremieux, Nicolas De Neuter, Benson Ogunjimi, Kris Laukens, Pieter Meysman

## Abstract

High-throughput T cell receptor (TCR) sequencing allows the characterization of an individual’s TCR repertoire and directly query their immune state. However, it remains a non-trivial task to couple these sequenced TCRs to their antigenic targets. In this paper, we present a novel strategy to annotate full TCR sequence repertoires. The strategy is based on a machine learning algorithm to learn the TCR patterns common to the recognition of a specific epitope. These results are then combined with a statistical analysis to evaluate the occurrence of specific epitope-reactive TCR sequences per epitope in repertoire data. In this manner, we can directly study the capacity of full TCR repertoires to target specific epitopes of the relevant vaccines or pathogens. We demonstrate the usability of this approach on three independent datasets related to vaccine monitoring and infectious disease diagnostics by independently identifying the epitopes that are targeted by the TCR repertoire. The developed method is freely available as a web tool for academic use at tcrex.biodatamining.be.

## INTRODUCTION

T cells form an important part of the adaptive immune system as they can recognize potentially pathogen-derived or aberrant peptides (epitopes), presented by major histocompatibility complex (MHC) molecules on the cell surface of nucleated host cells or professional antigen-presenting cells, and induce an immune response. The T cell receptor (TCR) molecule is responsible for the recognition of the MHC presented epitope. Each TCR protein is encoded by a genomic region that undergoes non-homologous recombination during T cell maturation, a process termed VDJ recombination. The randomness of the recombination process ensures the production of many different TCR proteins, with each T cell clone expressing a particular TCR, and allows the recognition of different epitopes. Epitope binding by the TCR is a critical step for the activation of targeted immune responses.

As high-throughput sequencing of TCR repertoires becomes more common, the need for more advanced analysis methods is increasing. A major challenge involves the functional identification of activated T cells from TCR repertoire data. To date, considerable efforts have been made to address this problem either by the detection of proliferated T cell clones, through the comparison of the abundance of TCR clonotypes between different time points (1, 2), or identification of enriched groups of similar T cell clones using VDJ recombination probabilities (3). Along with these identification methods, several strategies have been used to compare the TCR repertoires of pathology-associated tissues with healthy tissues or peripheral blood, for example in cancer (4, 5). However, the current identification methods lack the ability to discriminate disease-reactive T cells from other T cells that are playing a role in concurrent, non-related immune responses, thus requiring additional experiments. Different experimental strategies are currently available to determine epitope-specific TCRs, i.e. TCRs that recognize the epitope of interest, such as epitope-MHC multimer assays and peptide stimulation experiments. However, these techniques are not error-free as they can miss important epitope-specific TCRs (6, 7) or falsely report non-binding TCRs to be epitope specific (8). Nevertheless, these methods have been extensively used to generate a large amount of epitope-specific TCR data which has been collected in public databases such as the VDJdb (9) and McPAS-TCR (10).

Here, we present a new strategy to study T cell responses based on the direct identification of epitope-specific TCRs from TCR repertoire data. To determine the epitope specificity of TCRs, machine learning models have been developed to analyze TCR sequences and predict the probability that they recognize and bind specific epitopes. These methods are based on the principle that similar TCR sequences often target the same epitope (11) and that machine learning techniques can be used to learn the molecular underpinnings that are shared by these epitope-specific TCR sequences (12–14). While these methods have been shown to be performant on small targeted data sets, their application on full repertoire datasets remains challenging.

In this paper, we specifically address two issues that currently limit the use of prediction methods to full TCR repertoires. The first is the uncertainty on where to place the cut-off for a correct prediction. In other words, how confident must the prediction be before we accept it as truth. If it is chosen too strictly, we may miss many epitope-specific T cells. Conversely, if it is chosen too low, we increase the number of TCRs that are mistakenly identified as epitope-binding. To address this first issue, the strategy proposed here includes the estimation of a model-specific baseline prediction rate that can be used to set the prediction cut-off in an informed manner. The second issue is that the presence of epitope-specific TCRs is not restricted to TCR repertoires that have been stimulated by an epitope as they can also occur in small numbers in naïve, healthy TCR repertoires. Therefore, the identification of single epitope-specific TCR sequences by itself does not provide sufficient information on the capacity of the full TCR repertoire to target specific epitopes. Current prediction strategies do not evaluate whether a TCR repertoire as a whole contains more TCRs against a specific epitope than expected in a healthy TCR repertoire that is not enriched for these TCRs. To address this second issue, we introduce an enrichment analysis strategy to identify those specific epitopes that are being targeted within a given TCR repertoire.

We built upon the epitope-specific TCR classifier described in (12) and developed an approach designed for the identification of various epitope-specific TCRs in full repertoire datasets. We have applied this new approach to three independent datasets (1, 2, 15), which have previously been analyzed with the traditional TCR repertoire data analyses described above. In this manner, we show that by adding this layer of epitope information, we are able to link T cell data to broader immune repertoire targets. The described method is available as a web tool at tcrex.biodatamining.be.

## MATERIAL AND METHODS

### Collection of training data and assembly of the background TCR dataset

A positive training dataset was constructed containing human epitope-specific TCR beta chain sequences collected from the manually curated catalogue of pathology-associated T cell receptor sequences (McPAS-TCR (10)) (11 339 TCR-pathology combinations) and the VDJ database (VDJdb (9)) (17 792 TCR-epitope combinations) on November 18, 2018. To train the prediction models, both the CDR3 beta amino acid sequence and the V/J genes were gathered. The following quality filtering steps were applied for the McPAS-TCR dataset: (1) retaining epitope-specific TCRs determined with peptide-MHC tetramers or peptide stimulation and (2) removal of TCRs with missing information (i.e. CDR3 beta sequence, V/J genes or the specific epitope), reconstructed J genes, V/J genes with special characters that could not be matched to known V/J genes, CDR3 beta sequences with lower case amino acids or non-amino acid characters, TCRs with additional quality remarks and exclusion of TCRs from studies using mouse strains. The dataset was further extended with 550 TCR-epitope combinations found in the published literature (13, 16–18). Standard TCRex filtering steps were carried out (supplemental material S1). Furthermore, entries with multiple V genes were split into separate entries each having one of the V genes. For the retained TCR beta sequences, we checked whether their V/J genes were present in the IMGT format as formulated by the IMGT database (19). The latter was necessary due to possible inconsistencies in the way V/J genes were reported across studies. All V/J genes not reported in the IMGT format were corrected using our in-house IMGT parser. Finally, duplicate TCR-epitope combinations across different sources were removed. Only beta chain TCR sequence data for epitopes having at least 30 unique epitope-specific TCRs was retained. The final dataset contained 18 679 unique TCR-epitope combinations.

The control training dataset was designed to contain non-binding TCRs for each epitope. As one epitope can be presented by more than one MHC-molecule and one TCR sequence might interact with different MHC molecules (20), we did not take the epitopes’ MHC background into account. Instead, we assembled one control dataset for all epitope-specific models by collecting TCR beta chain sequences from healthy TCR repertoires from the study by Dean et al. (21). Although the vast majority of TCRs in healthy TCR repertoires will not target a certain epitope, it is not unlikely that these repertoires contain a few TCRs that do target the epitope of interest. The presence of these TCRs in our control dataset and their effect on the performance of the prediction models is expected to be negligible due to the low probability of their inclusion. In total, bulk TCR beta chain repertoire data from 587 healthy individuals was available. We randomly selected 1500 TCR beta sequences from the repertoire of each of these individuals. Subsequently standard TCRex filtering steps (supplemental material S1) were applied and the V/J genes of the remaining TCR beta chain sequences were transformed to the IMGT format. From this final dataset, 250 000 unique TCR beta chain sequences were randomly selected to make up the negative training dataset. Another different 100 000 unique TCR beta chain sequences were selected to represent the background dataset needed for the calculation of the baseline prediction rate.

### Collection of TCR datasets from infectious-disease studies

To evaluate the use of our epitope-specific models for the functional annotation of TCR repertoires, we collected TCR data from two independent yellow fever virus (YFV) vaccination studies (1, 2) and one cytomegalovirus (CMV) study (15). From the first YFV study (1), pre-(day 0) and post-vaccination (day 14) bulk TCR beta repertoire data (i.e. derived from peripheral blood mononuclear cells or PBMC) was collected from nine volunteers in the immunoSEQ Analyzer format. This study also reported the activated T cells in the post-vaccination repertoires for each of the volunteers (“CD3^+^ CD8^+^CD14^−^ CD19^−^CD38^+^HLA-DR^+^ Ag-experienced, activated effector T cells” (1)). From the second YFV study (2), pre-(day 0) and post-vaccination (day 15) bulk TCR repertoire data (i.e. derived from PBMC) was collected from three twins from https://github.com/mptouzel/pogorelyy_et_al_2018. In case biological replicates were present for a volunteer at a time point, these were merged. As the TCR repertoires from this study were reported in a MiXCR file format that did not follow the standard TCRex input requirements (i.e. two additional columns reporting the best V gene and the best J gene respectively for each TCR sequence), we parsed these files and selected the V/J gene with the highest score for every TCR sequence. For those TCRs with several V/J genes tied for the highest score, only the first one was retained. The second YFV study (2) also reported a list of vaccine-associated TCR beta sequences that were identified in the volunteers using their statistical analysis relying on clonal expansion. This list was used to compare the number of identified TCRs between TCRex and the YFV study. From the CMV study (15), we gathered a dataset of 164 TCRs that were reported to be CMV-associated following a statistical analysis of a large group (>600) of healthy CMV+ and CMV-volunteers.

### Model training and performance evaluation

For each epitope present in the collected training dataset, we trained a random forest model to identify epitope-binding TCRs in a TCR repertoire dataset. Training of epitope-specific prediction models was based on the method established in (12). This method was shown to perform comparably to other state-of-the-art classifiers in a recent independent study (22). In brief, the amino acid sequences of the CDR3 regions of the TCR beta chains were converted into physicochemical features and the beta chain’s V/J genes and families were one-hot encoded. For each epitope in the collected training data, a random forest classifier was trained with 100 individual decision trees using the scikit-learn library (23). We sampled 10 times more control training data than positive training data from the negative control set of 250 000 sequences to mimic the natural imbalance of epitope-specific T cells in full TCR repertoires. TCR beta chain sequences that occurred in both the positive training and negative training dataset were removed from the negative training dataset to avoid ambiguously labelled TCR beta chain sequences. Given a TCR beta chain sequence, each model returns the confidence value of binding a specific epitope. All models were evaluated using a stratified 5-fold cross-validation strategy. With this validation strategy we obtained the receiver operating characteristic (ROC) curve, precision-recall (PR) curve, the balanced accuracy, the area under the ROC curve (AUC) value and the average precision value for each classifier as implemented in the scikit-learn library (23). Models that had poor AUC (< 0.7) or poor average precision (< 0.35) values were excluded from the model collection.

### Controlling the baseline prediction rate

The class probability score for a TCR beta chain sequence is specific for each epitope model and does not take false positive predictions into account. To control the number of false positives, we provide a baseline prediction rate (BPR) value along with each sequence’s class probability score. This value represents the fraction of epitope-specific TCRs that can be detected in a background dataset (i.e. a control dataset representing a normal, healthy TCR repertoire) at a certain confidence cut-off. Since background datasets are expected to contain only a limited number of epitope-specific TCRs, this BPR can also be used as an estimate for the false positive rate (FPR). For every TCR sequence, its class probability score is compared to the class probability scores of the 100 000 unique background TCR beta chain sequences. A BPR value is then calculated for each TCR sequence as the fraction of randomly selected background sequences with a class probability score greater than or equal to the score of the TCR sequence. Thus, for a desired BPR threshold, a confidence cut-off *c* can be calculated so that:

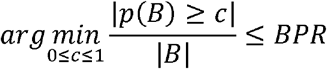

Where *p* is the prediction function and *B* is the background dataset. This allows us to filter with a custom threshold across different models by using for each the calculated cut-off *c*. Furthermore, this allows a direct interpretation of the cut-off with regards to the expected number of false positive hits for a typical TCR sequence dataset.

### Epitope specificity enrichment analysis

In addition to the detection of epitope-specific TCR beta chain sequences, we implemented a method to identify those epitopes for which we find a significantly higher number of TCR sequences in a target repertoire than expected in a representative control TCR repertoire. This analysis is performed by comparing the abundance of the epitope-specific TCR sequences in the TCR repertoire to a hypothetical dataset of randomly selected TCRs, i.e. the background dataset. This enrichment analysis strategy uses a BPR-based enrichment threshold that represents the percentage of identified epitope-specific TCRs in the background repertoire at a chosen BPR threshold. For each epitope of interest, a one-sided exact binomial test is used to compare the number of TCR sequences with a BPR score smaller or equal to the specified enrichment threshold. More specifically, we test whether we can reject the null hypothesis that the observed number of identified TCRs follows the binomial distribution *X~B*(*n, p*) and thus accept the alternative hypothesis that the abundance of TCRs specific for the epitope in the studied TCR repertoire exceeds the chosen enrichment threshold. Here, *X* is the number of identified TCRs specific for the studied epitope at a chosen enrichment threshold, *n* the size of the TCR repertoire and *p* the chosen enrichment threshold. The latter represents the percentage of epitope-specific TCRs that can be identified in a hypothetical background dataset at a specific BPR threshold. Since one false positive hit (i.e. a TCR sequence with a BPR score below the BPR threshold that does not recognize the epitope) is possible at every BPR threshold, we restrict the statistical analysis to those epitopes for which at least two epitope-specific TCR sequences are found.

### Web tool

The trained prediction models were collected and made available as a web application, called TCRex, which is accessible at tcrex.biodatamining.be. This web tool was built using the Django web framework (24) and makes use of several Python libraries including Scikit-learn (23), SciPy (25), Pyteomics (26), Altair (27), NumPy (28), Pandas (29) and Dask (30). It provides a user-friendly web interface to predict epitope binding for human TCR beta chain sequences and perform epitope specificity enrichment analyses for various viral and cancer epitopes. It is compatible with several common TCR sequence data formats, namely the immunoSEQ Analyzer formats (https://www.adaptivebiotech.com/immunoseq) and MiXCR (31) format, and the TCRex tab-delimited format which contains the V/J genes and CDR3 beta amino acid sequences for each TCR. More information concerning the use of the web tool is provided in supplemental material S2.

## RESULTS

### Overview of the epitope-specific models

The following performance metrics were calculated for each epitope-specific model using a 5-fold stratified cross-validation approach: the balanced accuracy, the area under the ROC curve (AUC) value and the average precision. These are shown in supplemental material S3. Only models with an AUC of at least 0.7 and an average precision of at least 0.35 were included in the final model collection, and thus in the current version (version 0.3.0) of the TCRex web tool. 49 out of 54 prediction models passed these inclusion criteria. Out of these 49 models, 44 were trained for viral epitopes and 5 for cancer epitopes (figure 1). The MHC-background for the epitopes is shown in supplemental material S4

**Figure 1:**
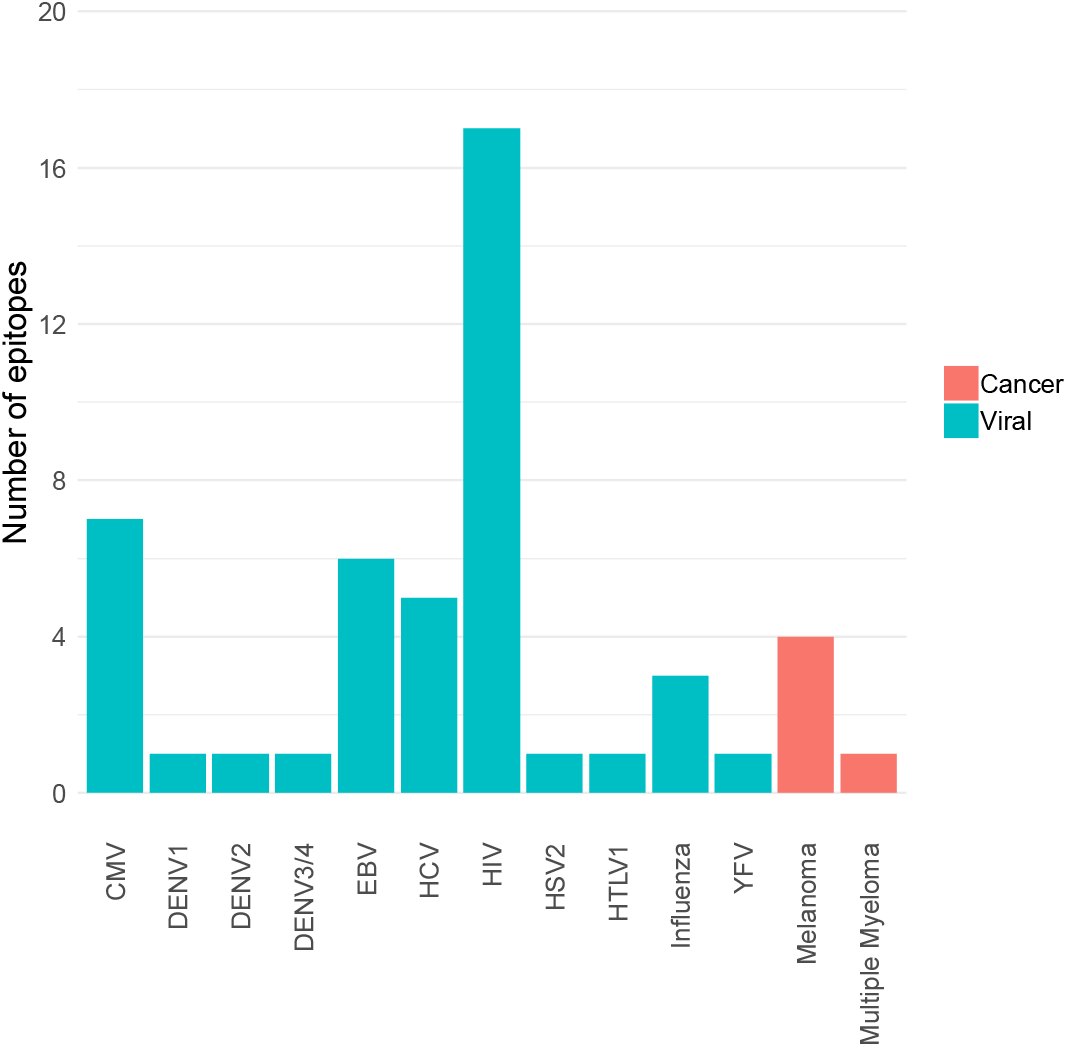
Overview of the number of trained prediction models for each virus and cancer type (TCRex version 0.3.0). The bars show the number of epitopes for which a prediction model with sufficient performance (i.e. AUC ≥ 0.7 and average precision ≥ 0.35) was trained.

### Evaluation of the prediction models using a leave-one-study-out validation approach

To evaluate how our epitope-specific models handle data from new studies, we designed a validation approach wherein all data of one study was left out during the training of the models and served as a holdout validation set. We selected viral epitopes with a large set of epitope-specific training data (at least 1000 unique TCRs) from various studies. For each of these epitopes, one study was removed from the training dataset (supplemental material S5). We ensured that this holdout dataset did not contain any of the training TCRs (i.e. TCRs with the same V/J genes and CDR3 beta sequence as training TCRs). After training the epitope-specific models, predictions on the holdout dataset were made using a strict BPR threshold of 0.01%. For each epitope-specific model, we expected a maximum of one false positive when analyzing these small datasets with a 0.01 % BPR threshold, as these datasets varied in size from 66 to 124 TCRs. To verify this assumption, the same models were used on a negative control dataset, i.e. a dataset lacking TCRs specific for the epitope of interest. For this purpose, we selected a set of TCRs recognizing the heteroclitic cancer epitope ELAGIGILTV. The number of ELAGIGILTV-specific TCRs was set to equal the size of the holdout validation datasets. We also confirmed that this negative test set did not contain any training TCRs, as this might be the case for putative cross-reactive TCRs. Standard performance metrics including accuracy, sensitivity, specificity and precision were calculated for each of the models (supplemental material S6). A summary of the results is shown in table 1. The majority of the identified TCRs did not match any of the training CDR3 beta sequences, which demonstrates the ability to find TCRs with unseen CDR3 beta sequences. Because of the strict BPR threshold, we achieved a very high specificity and precision. For both the GILGFVFTL and GLCTLVAML models, the specificity and precision reached the maximum score of 1 which corresponds to zero false positives. For the NLVPMVATV model, two false positives were detected, which does not align with the chosen BPR threshold. One of these TCRs, however, shares its CDR3 beta sequence and J gene with a NLVPMVATV-specific training TCR. This may indicate the presence of cross-reactive TCR recognition patterns for both the cancer and the CMV epitope. If this is the case, the TCR could be considered a true positive. The presence of epitope-specific TCRs in background datasets is a potential issue, as epitope-specificity studies mainly report TCRs binding to the epitope of interest and not the non-binding TCRs. Since there is no public information available on non-binding TCRs, our control datasets might contain a small number of epitope-specific TCRs. This could have led to an overestimation of the number of false positives in this analysis. Overall, the models performed well and did not seem to be subject to any study specific biases.

**Table 1:** Validation of three epitope-specific models with the leave-one-study-out validation approach. For each epitope, the size of the training dataset, the size of the test datasets (i.e. the holdout validation dataset and the additional cancer dataset) and the number of identified epitope-specific TCRs in the test datasets is given together with the associated performance metrics for a BPR threshold of 0.01%. In addition, the number of identified TCRs in the holdout validation set having a training CDR3 beta sequence is shown.

| Epitope | Size training dataset | Size test datasets | Identified TCRs in epitope-specific test dataset | Identified TCRs containing training CDR3 beta sequences | Identified TCRs in cancer dataset | Performance metrics |  |  |  |
| --- | --- | --- | --- | --- | --- | --- | --- | --- | --- |
|  |  |  |  |  |  | Accuracy | Sensitivity | Specificity | Precision |
| NLVPMVATV | 4723 | 89 | 10 | 3 | 2 | 0.54 | 0.11 | 0.98 | 0.83 |
| GILGFVFTL | 2983 | 124 | 3 | 0 | 0 | 0.51 | 0.02 | 1 | 1 |
| GLCTLVAML | 1142 | 66 | 12 | 3 | 0 | 0.59 | 0.18 | 1 | 1 |

### The number of unique epitope-specific TCRs increases post-vaccination

Previous studies have already demonstrated the feasibility of identifying T cell clones that have proliferated between time points in TCR repertoire data (1, 2). However, such methods do not provide insight into the targeted epitopes. Considering the fact that T cells proliferate upon binding with vaccine-associated epitopes, we hypothesized that post-vaccination repertoires contain an elevated level of unique epitope-specific TCRs compared to pre-vaccination repertoires. To evaluate the difference in the number of epitope-specific TCRs captured within the sequenced repertoire before and after vaccination, we reanalyzed published TCR repertoires from two independent YFV vaccination studies. In the first study, nine healthy volunteers with unknown HLA types received the YF-17D vaccine (1). In the second study, the YF-17D vaccine was used to vaccinate three HLA-A*02:01 positive twins (2). For each of the participants, TCR beta repertoire data from PBMC samples taken before and two weeks after vaccination were available. At the time of analysis, TCRex featured one YFV epitope prediction model: the immunodominant LLWNGPMAV peptide. We used this model to identify LLWNGPMAV-specific TCRs in both pre- and post-vaccination repertoires. To control the number of false positives, a strict BPR threshold of 0.01% was applied. As can be seen in figures 2A and 2B, the percentage of unique LLWNGPMAV-specific TCR sequences increased post vaccination for the majority of the volunteers (one-sided Wilcoxon signed-rank test, p value = 0.03711 for (1) and p value = 0.01563 for(2)).

**Figure 2:**
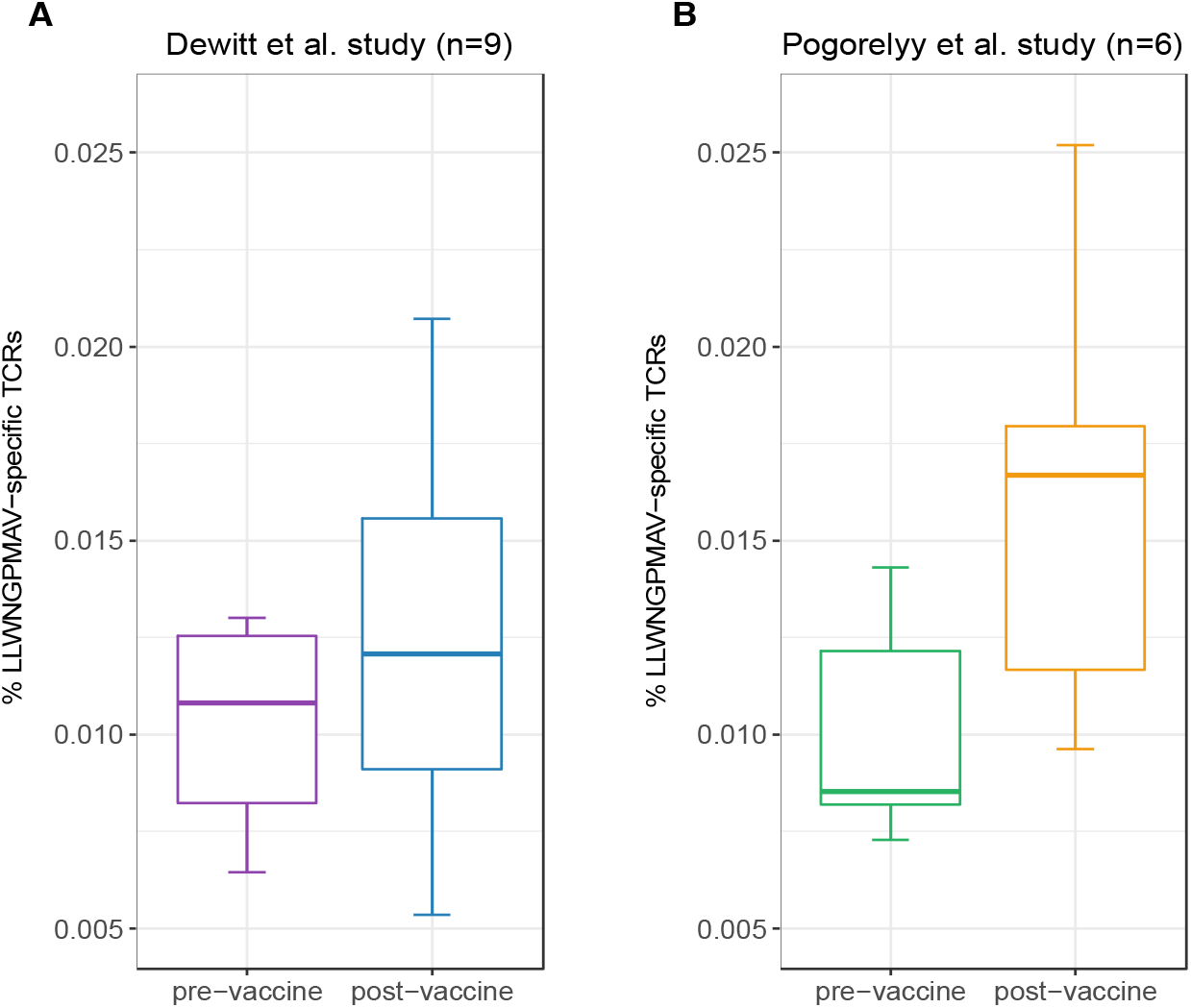
Percentage of identified LLWNGPMAV-specific TCRs pre- and post-vaccination. The boxplots show the proportion of unique LLWNGPMAV-specific T cells in pre- and post-vaccination PBMC samples for the nine volunteers from (1) (A) and the six volunteers from (2) (B). Epitope-specific T cells were identified with TCRex using a 0.01% BPR threshold. An increase in the number of unique epitope-specific cells was found for both studies.

### Comparison of the identification level with techniques relying on clonal expansion

The detection of vaccine-associated T cells with our new method is restricted to the recognition of epitope-specific T cells for which trained prediction models are available. Because only one YFV epitope model passed the quality filters at the time of analysis, we expected to detect a considerably smaller number of vaccine-associated TCRs in comparison with a differential analysis between time points. To evaluate the identification level of our new strategy, we compared our results from the post-vaccination repertoires to the outcome of the original YFV vaccine study from (2) which reported a list of the significantly expanded TCR clones two weeks after vaccination. Two important findings resulted from this analysis. First, we demonstrate that method (2) detected between 3 and 17 times the number of TCRs than TCRex. Second, the overlap between the epitope-specific TCRs identified by TCRex and the proliferated TCRs identified by (2) is rather limited, with an overlap ranging from 2 to 19 TCRs (table 2). These results could indicate that our approach might be able to detect new epitope-specific TCRs that might be missed by other techniques. The expanded TCR clones we could not identify were likely derived from T cells reacting with other YFV epitopes. Additionally, as the clonal expansion techniques do not take into account the epitope specificity of the TCRs, they might also falsely report bystander TCRs or TCRs associated with other non-vaccine related immune responses. Indeed, in the original YFV vaccine study three different experimental assays (i.e. a multimer assay using the LLWNGPMAV epitope, evaluation of the interferon gamma production following vaccine-stimulation and identification of activated CD8^+^CD38^+^HLA–DR^+^ T cells) were carried out to assess whether the identified expanded T cell clones for one of the volunteers (S1) were vaccine-associated (2). About half of the expanded 773 TCR clones were experimentally validated, of which 68 clonotypes were found to be LLWNGPMAV-specific. This is less than the 170 epitope-specific TCRs identified with TCRex. Although tetramer assays are known to miss epitope-specific TCRs (6, 7), this low identification rate might also be a consequence of the fact that the multimer assay was performed two years after vaccination.

**Table 2:** Comparison of the epitope-specific TCR sequences identified with TCRex to the results from (2). For each volunteer from (2) are given: the number of YFV-specific clones (DNA level) reported by the YFV vaccine study, the number of unique canonical YFV-specific TCRs (i.e. TCRs starting with cysteine and/or ending with phenylalanine) (protein level with CDR3 beta sequences not containing any non-amino acid characters) identified in the YFV vaccine study, the number of epitope-specific sequences in the post-vaccination repertoire identified by TCRex and the number of TCRs identified by both the YFV vaccine study and TCRex.

| Volunteer | Vaccine-associated TCR clones reported by (2) | Unique canonical vaccine-associated TCRs identified by (2) | Epitope-specific TCRs identified by TCRex | Overlapping TCRs |
| --- | --- | --- | --- | --- |
| P1 | 1151 | 1090 | 92 | 2 |
| P2 | 800 | 763 | 110 | 3 |
| Q1 | 576 | 547 | 79 | 7 |
| Q2 | 1685 | 1589 | 92 | 8 |
| S1 | 773 | 747 | 170 | 19 |
| S2 | 983 | 948 | 273 | 11 |

### The majority of the identified LLWNGPMAV-specific T cells are not present in public databases

Since the detection of epitope-specific T cells relies on a prediction model trained on previously identified TCRs recognizing the epitope, we assessed to what extent our models are able to identify new epitope-specific TCRs. For this, we compared the number of TCRex-identified LLWNGPMAV-specific T cells in the post-vaccination repertoires from the two YFV vaccine studies (1, 2) with the number of public TCRs (i.e. known LLWNGPMAV-specific TCR sequences present in the training data) in these repertoires. For both YFV studies, TCRex achieved a higher identification rate in comparison with the database search technique (figure 3). The number of TCRs identified by both techniques is rather limited (supplemental material S7). TCRex does find some training TCRs in the repertoires, however, we can deduce that the bulk of the TCRs identified by TCRex were not present in our training data and thus not publicly known.

**Figure 3:**
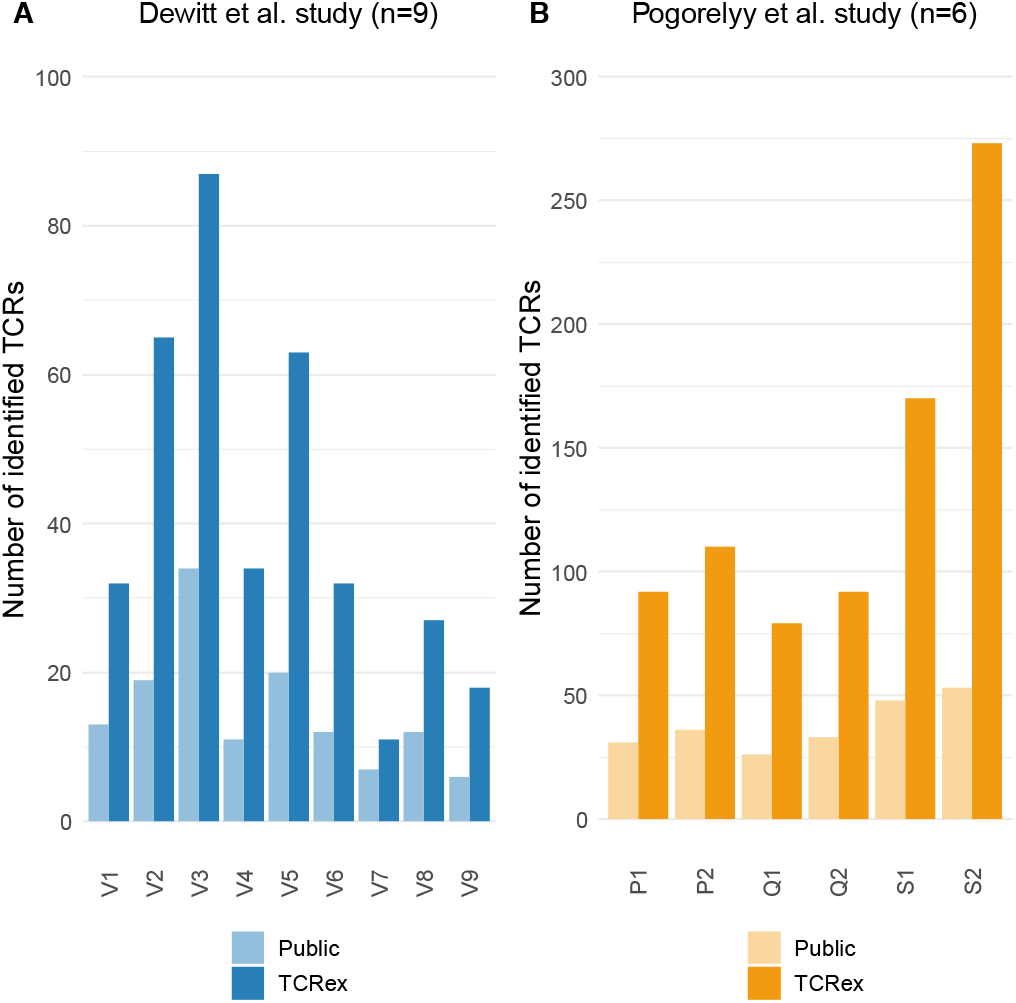
Public versus TCRex-identified LLWNGPMAV-specific TCRs in the post-vaccination samples of (1) and (2). (A) Overview of the number of unique LLWNGPMAV-specific TCRs that were identified with TCRex in the post-vaccination PBMC samples for all nine volunteers (V1-V9) of (1) (dark blue) and the total number of public LLWNGPMAV-specific TCRs that were present in these repertoires (light blue). (B) Overview of the number of unique LLWNGPMAV-specific TCRs that were identified with TCRex in the post-vaccination PBMC samples for all six volunteers (P1-S2) of (2) (dark orange) and the total number of public LLWNGPMAV-specific TCRs that were present in these repertoires (light orange).

### Identification of enriched vaccine-associated TCR repertoire targets

In addition to the identification of epitope-specific TCRs in whole TCR repertoires, our newly developed strategy can also be used to detect enriched TCR repertoire targets. In essence, we detect epitopes for which more specific TCRs are present than expected based on the epitope-specific TCR occurrence in a healthy, non-vaccinated TCR repertoire. Here, we demonstrate the feasibility of this enrichment analysis by comparing the levels of LLWNGPMAV-specific TCRs before and after vaccination for the volunteers from the two YFV studies. For each of the volunteers, we evaluated whether their pre- and post-vaccination TCR repertoires contained more LLWNGPMAV-specific TCRs than expected in a background TCR repertoire, i.e. a TCR dataset representing a healthy, non-vaccinated TCR repertoire. As all the LLWNGPMAV-specific prediction analyses were performed with a 0.01% BPR threshold, we expect that the abundance of LLWNGPMAV-specific TCRs in a background repertoire will not exceed 0.01%. Therefore, we assessed for each of the volunteers’ TCR repertoires whether the percentage of epitope-specific TCRs in the TCR repertoires was significantly higher than 0.01% and thus were enriched in LLWNGPMAV-specific TCRs. The number of identified TCRs and the results of the epitope-specificity enrichment analyses for all volunteers from (2) are shown in table 3. The majority of the post-vaccination TCR repertoires (4 out of 6) are enriched in LLWNGPMAV-specific TCRs, as opposed to the pre-vaccination TCR repertoires (2 out of 6). The enrichment in pre-vaccination repertoires might be explained by the use of an unrepresentative background dataset, which is a common problem addressed in the next section. A similar trend can be seen for the volunteers from (1). Here, the post-vaccination TCR repertoires from 3 of the 9 volunteers demonstrated an elevated level of LLWNGPMAV-specific TCRs while no volunteer showed an enrichment in their pre-vaccination TCR repertoire (table 4).

**Table 3:** LLWNGPMAV-specific enrichment analysis results for all volunteers from (2). The table represents the q values (Benjamini-Hochberg corrected p values for 6 tests) associated with the LLWNGPMAV-specific enrichment analyses for both full (i.e. derived from PBMC) pre- and post-vaccination TCR repertoires. In addition, the number of identified specific TCRs and the percentage of identified specific TCRs are given.

| Volunteer | Identification level<br>pre-vaccination<br>PBMC |  |  | Identification level<br>post-vaccination<br>PBMC |  |  |  |
| --- | --- | --- | --- | --- | --- | --- | --- |
|  | Identified<br>TCRs | %<br>epitope-specific<br>TCRs | q value<br>enrichment<br>analysis vs.<br>background<br>repertoire | Identified<br>TCRs | %<br>epitope-specific<br>TCRs | q value<br>enrichment<br>analysis vs.<br>background<br>repertoire | q value<br>enrichment<br>analysis<br>vs. pre-vaccination<br>repertoire |
| P1 | 77 | 0.007272 | 0.999 | 92 | 0.010400 | 0.441 | 8.17e-04 |
| P2 | 95 | 0.008119 | 0.999 | 110 | 0.009614 | 0.673 | 0.045 |
| Q1 | 61 | 0.008615 | 0.999 | 79 | 0.017915 | 2.80e-06 | 1.24e-08 |
| Q2 | 109 | 0.008440 | 0.999 | 92 | 0.015455 | 8.59e-05 | 1.64e-07 |
| S1 | 138 | 0.014311 | 2.40e-04 | 170 | 0.025194 | 6.06e-25 | 5.69e-11 |
| S2 | 180 | 0.013314 | 4.09e-04 | 273 | 0.017962 | 2.10e-18 | 2.03e-06 |

**Table 4:** LLWNGPMAV-specific enrichment analysis results for all volunteers from (1). The table represents the q values (Benjamini-Hochberg corrected p values for 9 tests) associated with the LLWNGPMAV-specific enrichment analyses for both full (i.e. derived from PBMC) pre- and post-vaccination TCR repertoires and the post-vaccination activated TCR repertoire. In addition, the number of identified specific TCRs and the percentage of identified specific TCRs are given.

| Volunteer | Pre-vaccination<br>full TCR repertoire |  |  | Post-vaccination<br>full TCR repertoire |  |  |  | Post-vaccination<br>activated TCR repertoire |  |  |  |
| --- | --- | --- | --- | --- | --- | --- | --- | --- | --- | --- | --- |
|  | Identified<br>TCRs | %<br>epitope-specific<br>TCRs | q value<br>enrichment<br>analysis<br>vs.<br>background<br>repertoire | Identified<br>TCRs | %<br>epitope-specific<br>TCRs | q value<br>enrichment<br>analysis<br>vs.<br>background<br>repertoire | q value<br>enrichment<br>analysis<br>vs. pre-vaccination<br>repertoire | Identified<br>TCRs | %<br>epitope-specific<br>TCRs | q value<br>enrichment<br>analysis<br>vs.<br>background<br>repertoire | q value<br>enrichment<br>analysis<br>vs. pre-vaccination<br>repertoire |
| 1 | 35 | 0.010813 | 0.621 | 32 | 0.009094 | 0.935 | 0.857 | 4 | 0.017189 | 0.265 | 0.316 |
| 2 | 48 | 0.012999 | 0.384 | 65 | 0.015565 | 1.56e-03 | 0.212 | 5 | 0.028991 | 0.056 | 0.116 |
| 3 | 41 | 0.012539 | 0.384 | 87 | 0.025967 | 9.29e-14 | 7.86e-09 | 181 | 0.550922 | 1.86e-239 | 5.00e-222 |
| 4 | 23 | 0.009778 | 0.856 | 34 | 0.012084 | 0.299 | 0.231 | 21 | 0.103128 | 2.56e-14 | 1.67e-14 |
| 5 | 8 | 0.012759 | 0.621 | 63 | 0.020710 | 7.22e-07 | 9.93e-04 | 71 | 0.423527 | 7.50e-87 | 1.56e-79 |
| 6 | 41 | 0.012111 | 0.384 | 32 | 0.012065 | 0.299 | 0.684 | 8 | 0.040422 | 2.31e-03 | 7.45e-03 |
| 7 | 11 | 0.006771 | 0.964 | 11 | 0.005344 | 0.992 | 0.857 | 2 | 0.011544 | 0.517 | 0.369 |
| 8 | 22 | 0.008229 | 0.964 | 27 | 0.010866 | 0.539 | 0.212 | 2 | 0.011869 | 0.517 | 0.404 |
| 9 | 13 | 0.006451 | 0.964 | 18 | 0.008163 | 0.938 | 0.284 | 4 | 0.021606 | 0.176 | 0.060 |

For the YFV study from (1) we additionally studied the abundance of LLWNGPMAV-specific TCRs in experimentally enriched TCR repertoires as Dewitt et al. reported the activated subset (i.e. “CD3^+^ CD8^+^CD14^−^ CD19^−^CD38^+^HLA-DR^+^ Ag-experienced, activated effector T cells” (1)) of the TCR repertoire for each of the nine volunteers 14 days after vaccination with the YFV-17D vaccine. For each of these activated TCR repertoires, LLWNGPMAV-specific TCRs were identified using a BPR threshold of 0.01% and compared with the abundance of specific TCRs in the unsorted PBMC samples at the same time point. The number of LLWNGPMAV-specific TCRs in the activated TCR dataset increased drastically in comparison to their unsorted PBMC samples for the majority of the volunteers (one-sided Wilcoxon signed-rank test, p value = 0.001953) (figure 4). The same trend can be seen in the results of the enrichment analysis, presented in table 4, with 4 out of 9 of the activated TCR repertoires demonstrating an enrichment in LLWNGPMAV-specific TCRs whereas this is limited to 3 out of 9 for the unsorted post-vaccination repertoires.

**Figure 4:**
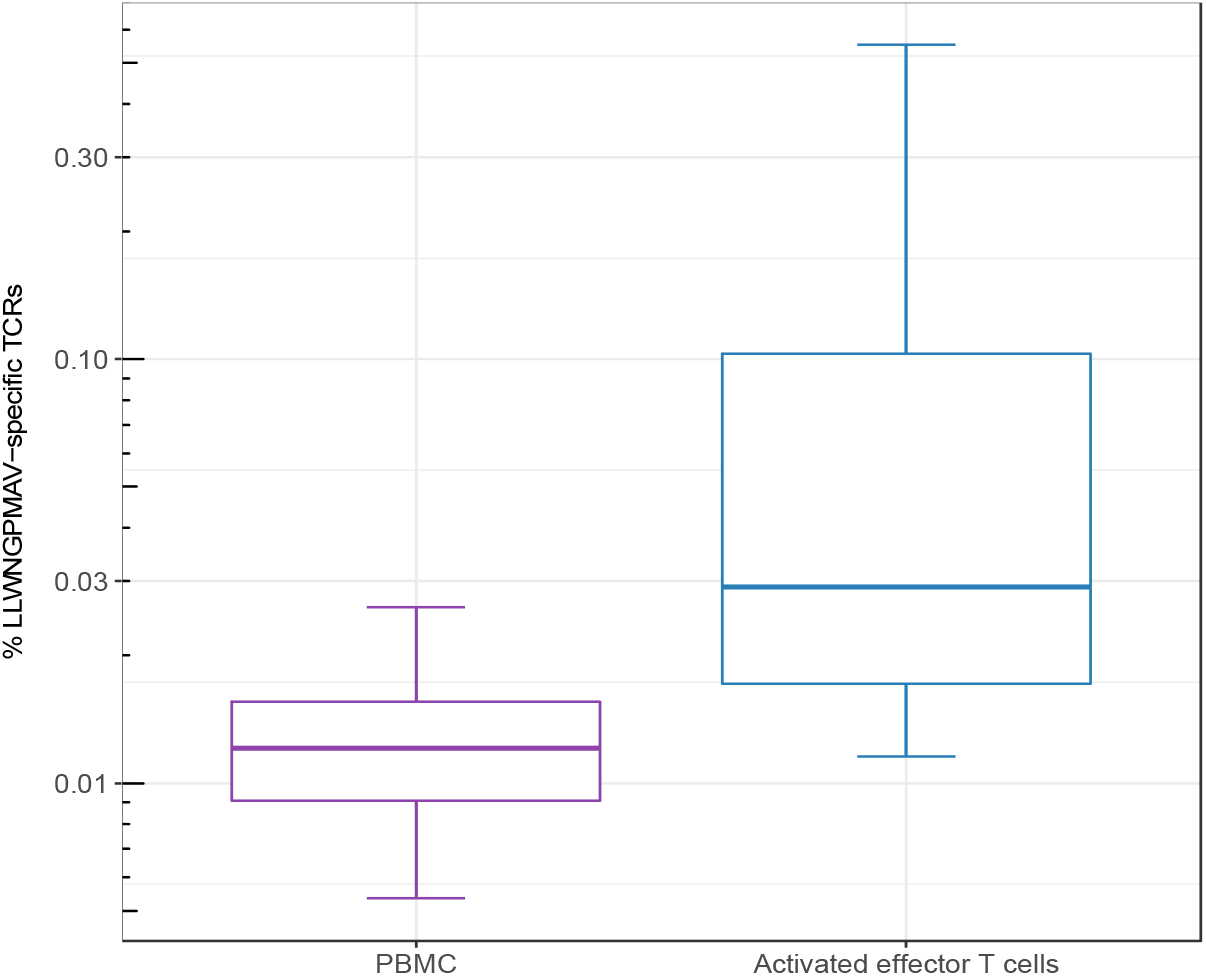
Percentage of identified LLWNGPMAV-specific TCRs in the post-vaccination repertoires from (1). The box plots show the log scaled proportion of unique LLWNGPMAV-specific T cells in post-vaccination PBMC samples and activated TCR repertoires (i.e. “CD3+ CD8+CD14-CD19-CD38+HLA-DR+ Ag-experienced, activated effector T cells” (1)) for the nine volunteers from (1). Epitope-specific TCRs were identified with TCRex using a 0.01% BPR threshold. An increase in the number of unique LLWNGPMAV-specific cells was found in the activated dataset.

### The importance of the background dataset in enrichment analyses

In the previous section, we showed how an enrichment in epitope-specificity in TCR repertoires can be detected by comparing the abundances of epitope-specific TCRs to a general background dataset with a predefined enrichment threshold. Here, we demonstrate how the selection of a representative background dataset can improve these enrichment analyses. For each of the volunteers from the two YFV studies (1, 2), we compared the abundance of LLWNGPMAV-specific TCRs in their post-vaccination repertoire to the abundance in their pre-vaccination repertoire. For this, we slightly adapted our general epitope specificity enrichment analysis: (1) the number of successes in the one-sided exact binomial test was set equal to the number of epitope-specific TCRs in the post-vaccination repertoire that was identified with a 0.01% BPR threshold and (2) the enrichment threshold was set equal to the fraction of the number of identified TCRs in their pre-vaccination repertoire. A comparison between the two different enrichment analysis strategies demonstrates the importance of the use of a representative background dataset (tables 3 and 4). When the percentage of epitope-specific TCRs in the pre-vaccination repertoires is much lower than we generally expect in healthy TCR repertoires, the use of a standard background dataset might not discover the increase of epitope-specific TCRs in post-vaccination repertoires. This can be seen for volunteer P1 and P2 from study (2) (table 3). Conversely, when the percentage of epitope-specific TCRs in the pre-vaccination repertoires exceeds our expectations, the use of a background dataset could falsely indicate a significant increase of epitope-specific TCRs in post-vaccination repertoires. This can be seen for volunteer 2 from study (1) (table 4). In conclusion, the selection of the background dataset is an important step when performing enrichment analyses and depends on the research question and experimental set-up of the study.

### Identification of enriched disease-associated TCR repertoire targets

Previously, Emerson et al. introduced a new statistical method to discover disease-associated TCRs from TCR repertoire data (15). Briefly, they sequenced the TCR beta repertoires of a large group (more than 600) healthy CMV^+^ and CMV^−^ volunteers and set up a statistical test to identify public TCRs that were enriched in the healthy CMV^+^ volunteers. In total, 164 TCRs were found to be associated with a positive cytomegalovirus serostatus of which three were matched with a CMV epitope through an additional validation experiment. Here, we use our enrichment analysis strategy to identify the CMV epitopes that are targeted by the TCRs in this dataset. In short, we searched for epitopes for which an elevated level of specific TCRs are present in this dataset, i.e. more than expected in a background TCR repertoire. To see whether we only find an enrichment for CMV-specific TCR sequences, and not for non-CMV epitopes, we performed binding predictions for all epitopes present in TCRex with a strict BPR threshold of 0.01%. In total, 8 out of 164 different TCRs were predicted to bind to one of the CMV epitopes (table 5). Two epitopes, both derived from CMV, were found to be significantly enriched in the dataset: NLVPMVATV (q value= 7.1e-7, Benjamini-Hochberg corrected p value for two tests) and TPRVTGGGAM (q value= 5.7e-09, Benjamini-Hochberg corrected p value for two tests), compared to a TCR background dataset where we would expect to identify 0.01% specific TCRs for each epitope with our selected BPR threshold. Additionally, we also found one YFV-specific TCR in the list of 164 CMV associated TCR sequences. This might be a false positive identification, where the LLWNGPMAV-specific model wrongly classified the TCR as epitope-binding. However, this TCR sequence shared an identical combination of a CDR3 beta sequence and J gene with TCR sequences linked with the YFV epitope in public databases and thus the model training set. It is therefore possible that this TCR is cross-reactive and recognizes both the immunodominant YFV epitope and a CMV epitope. It could also be that the identified TCR is not CMV-reactive as the dataset reported by the original study represents TCR sequences that were statistically associated with the CMV seropositive volunteers which does not directly entail CMV-reactivity. In summary, this analysis indicates the relevance of using prediction models to identify epitope-specific T cells and search for enriched T cell epitopes in a TCR beta dataset.

**Table 5:**
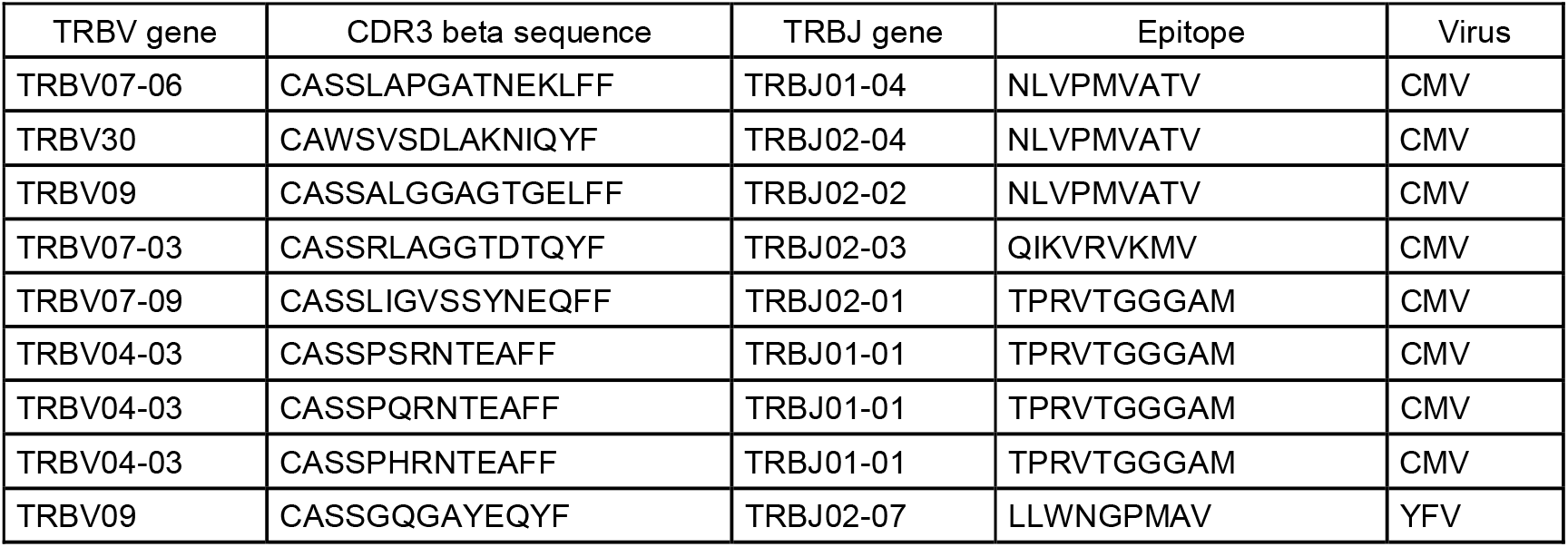
Epitope-specific TCRs identified in the enriched CMV-related TCR dataset of (15) (0.01% BPR threshold).

## DISCUSSION

Traditional methods to identify epitope-specific T cells require targeted experiments, which are known to fail to identify a considerable amount of reactive T cells (6, 7). Approaches to solve this problem computationally have been proposed, but they are hard to translate to full repertoires due to their inherent prediction error. To overcome these limitations, we propose a novel approach built on the identification of epitope-specific T cells. Our method uses epitope-specific prediction models trained on publicly available TCR beta sequence data to assess the probability of binding the epitope of interest and corrects for prediction errors by using a stringent BPR threshold. As such, it allows for the easy identification of epitope-specific T cells in full TCR repertoires and consequently the detection of enriched TCR repertoire targets. In the current implementation, it is based on a random forest classifier trained on common physicochemical properties. The implemented method currently only uses TCR beta chain information, as this is the bulk of the data that is currently available. As linked beta-alpha TCR data becomes more common (32, 33), the approach can be easily extended to include alpha chain sequence data.

This novel approach was validated on several independent datasets to address different research questions. Firstly, we examined whether we could detect a difference in the presence of epitope-specific TCRs in pre- and post-vaccination repertoires. Considering that vaccines are expected to activate T cells upon binding with vaccine-associated epitopes, leading to a targeted proliferation of these T cells, we expected to find more unique epitope-specific T cells after vaccination. The analysis of the two YFV studies (1, 2) seems to support this hypothesis as an increase in LLWNGPMAV-specific T cells is found in the post-vaccination repertoires from both studies. In comparison with the method of (2), we achieve a relatively high identification rate considering our analysis was restricted to one epitope. However, the majority of the TCRs identified with TCRex do not overlap with the results in (2). This could indicate that we might be able to find epitope-specific TCRs that are missed by other methods. For example, it is possible that some epitope-specific TCRs expanded less in comparison to others and were therefore not detected using the technique relying on clonal expansion. Though more extensive experimental validation would be required, TCRex has the potential to be used as a complementary technique to existing TCR data analysis strategies by annotating single TCRs with their epitope target and/or by discovering epitope response trends in (activated) TCR repertoires.

Finally, we demonstrated how our approach enables the search for enriched epitopes in TCR repertoire datasets using our built-in enrichment analysis test. In short, we used an enrichment test to check whether a TCR repertoire contains significantly more TCRs recognizing a specific epitope than a random background TCR dataset. As a proof of concept, we compared the elevated level of LLWNGPMAV-specific TCRs in pre- and post-vaccination TCR repertoires and analyzed a dataset enriched in CMV-associated TCRs. The former analysis shows the importance of the selection of the right background dataset when performing enrichment analyses, which we discuss in more detail in the next section. For the latter, our enrichment analysis method easily detected a significant increase in epitope-specific TCRs for two CMV epitopes. No enrichment was found for other viral epitopes, which illustrates the robustness of our approach.

While this novel approach allows a full repertoire exploration, several potential sources of bias may still impact the results. For example, it is unclear to what extent epitope-specific TCRs are also MHC-restricted and thus only bind a certain epitope when it is presented on a specific MHC. Since our epitope-specific models were trained on TCR sequence data with a restricted MHC background, they might be specific for the epitope-MHC complex. This potential bias could lead to a bias in the enrichment results when studying the TCR repertoires of individuals whose HLA types do not match the MHC background of our prediction models as epitope-specific TCRs with other MHC-specificities might not be detected. However, we do have some reasons to believe that the epitope-specific prediction models are MHC-agnostic. First, previous studies have shown that TCRs can be shared across individuals with different HLA-types (20) and that epitope-specific TCR patterns seem to transcend the MHC background of the epitopes (11). This might indicate that epitope-specific TCR sequences can recognize a specific epitope even when it is presented by different MHCs. Second, we demonstrated that we can identify enriched epitope-specificities in the CMV-associated TCR dataset which was assembled from multiple CMV seropositive individuals having different HLA types. This suggests that our method can work on TCR datasets with different MHC backgrounds. However, even if the epitope-specific models were not MHC-agnostic, potential biases in the enrichment results can be avoided by comparing the abundance of epitope-specific TCRs in a TCR repertoire to a background dataset with a matching MHC background. In general, we recommend the use of a representative background dataset (e.g. background data from the same subject using the same TCR sequencing and analysis set up) when available as any other biases between the TCR repertoire and the background dataset might affect the enrichment results. This is demonstrated in the enrichment analyses of the YFV studies where the replacement of the standard background dataset by the pre-vaccination repertoires gave better results. Nevertheless, if no suitable background dataset is available, the enrichment analysis, as implemented in the TCRex web tool, still provides an easy and fast exploration of a TCR dataset, since it gives an initial overview of the possible TCR targets that may be enriched in this dataset. As bulk sequenced TCR repertoires are commonly not annotated for any epitope target and require laborious additional experiments to discover these targets, this enrichment analysis can inform subsequent experimental efforts.

Taken together, we have developed a novel strategy to study the epitope-specificity of full TCR repertoires. Our approach relies on the identification of epitope-specific T cell receptor sequences using epitope-specific prediction models. This enables the prediction of human epitope-specific TCR sequences and the identification of enriched TCR repertoire targets. We have embedded our method into a user-friendly web tool, called TCRex, which is available at tcrex.biodatamining.be. This tool will facilitate the study of the epitope specificity of TCR sequences that underly immune responses in different disease-associated studies.

## REFERENCES

1. Dewitt,W.S., Emerson,R.O., Lindau,P., Vignali,M., Snyder,T.M., Desmarais,C., Sanders,C., Utsugi,H., Warren,E.H., McElrath,J., et al. (2015) Dynamics of the Cytotoxic T Cell Response to a Model of Acute Viral Infection. J. Virol., 89, 4517–4526.

2. Pogorelyy,M. V., Minervina,A.A., Touzel,M.P., Sycheva,A.L., Komech,E.A., Kovalenko,E.I., Karganova,G.G., Egorov,E.S., Komkov,A.Y., Chudakov,D.M., et al. (2018) Precise tracking of vaccine-responding T cell clones reveals convergent and personalized response in identical twins. PNAS, 115, 12704–12709.

3. Pogorelyy,M. V, Minervina,A.A., Shugay,M., Chudakov,D.M., Lebedev,Y.B., Mora,T. and Walczak,A.M. (2019) Detecting T-cell receptors involved in immune responses from single repertoire snapshots. PLoS Biol., 17, e3000314.

4. Sherwood,A., Emerson,R., Scherer,D., Habermann,N., Buck,K., Staffa,J., Desmarais,C., Halama,N., Jaeger,D., Schirmacher,P., et al. (2013) Tumor-infiltrating lymphocytes in colorectal tumors display a diversity of T cell receptor sequences that differ from the T cells in adjacent mucosal tissue. Cancer Immunol. Immunother., 62, 1453–1461.

5. Ahmadzadeh,M., Pasetto,A., Jia,L., Deniger,D.C., Stevanović,S., Robbins,P.F. and Rosenberg,S.A. (2019) Tumor-infiltrating human CD4 + regulatory T cells display a distinct TCR repertoire and exhibit tumor and neoantigen reactivity. Sci. Immunol., 4, eaao4310.

6. Rius,C., Attaf,M., Tungatt,K., Bianchi,V., Legut,M., Bovay,A., Donia,M., Thor Straten,P., Peakman,M., Svane,I.M., et al. (2018) Peptide–MHC Class I Tetramers Can Fail To Detect Relevant Functional T Cell Clonotypes and Underestimate Antigen-Reactive T Cell Populations. J. Immunol., 200, 2263–2279.

7. Dolton,G., Zervoudi,E., Rius,C., Wall,A., Thomas,H.L., Fuller,A., Yeo,L., Legut,M., Wheeler,S., Meriem,A., et al. (2018) Optimized Peptide –MHC Multimer Protocols for Detection and Isolation of Autoimmune T-Cells. Front. Immunol., 9, 1378.

8. Martin,M.D., Jensen,I.J., Ishizuka,A.S., Lefebvre,M., Shan,Q., Xue,H.-H., Harty,J.T., Seder,R.A. and Badovinac,V.P. (2019) Bystander responses impact accurate detection of murine and human antigen-specific CD8 T cells. J. Clin. Invest., 10.1172/JCI124443.

9. Shugay,M., Bagaev,D. V, Zvyagin,I. V, Vroomans,R.M., Crawford,J.C., Dolton,G., Komech,E.A., Sycheva,A.L., Koneva,A.E., Egorov,E.S., et al. (2018) VDJdb: a curated database of T-cell receptor sequences with known antigen specificity. Nucleic Acids Res., 46, D419–D427.

10. Tickotsky,N., Sagiv,T., Prilusky,J., Shifrut,E. and Friedman,N. (2017) McPAS-TCR: a manually curated catalogue of pathology-associated T cell receptor sequences. Bioinformatics, 33, 2924–2929.

11. Meysman,P., De Neuter,N., Gielis,S., Bui Thi,D., Ogunjimi,B. and Laukens,K. (2019) On the viability of unsupervised T-cell receptor sequence clustering for epitope preference. Bioinformatics, 35, 1461–1468.

12. De Neuter,N., Bittremieux,W., Beirnaert,C., Cuypers,B., Mrzic,A., Moris,P., Suls,A., Van Tendeloo,V., Ogunjimi,B., Laukens,K., et al. (2018) On the feasibility of mining CD8+ T cell receptor patterns underlying immunogenic peptide recognition. Immunogenetics, 70, 159–168.

13. Dash,P., Fiore-Gartland,A.J., Hertz,T., Wang,G.C., Sharma,S., Souquette,A., Crawford,J.C., Clemens,E.B., Nguyen,T.H.O., Kedzierska,K., et al. (2017) Quantifiable predictive features define epitope-specific T cell receptor repertoires. Nature, 547, 89–93.

14. Glanville,J., Huang,H., Nau,A., Hatton,O., Wagar,L.E., Rubelt,F., Ji,X., Han,A., Krams,S.M., Pettus,C., et al. (2017) Identifying specificity groups in the T cell receptor repertoire. Nature, 547, 94–98.

15. Emerson,R.O., DeWitt,W.S., Vignali,M., Gravley,J., Hu,J.K., Osborne,E.J., Desmarais,C., Klinger,M., Carlson,C.S., Hansen,J.A., et al. (2017) Immunosequencing identifies signatures of cytomegalovirus exposure history and HLA-mediated effects on the T cell repertoire. Nat. Genet., 49, 659–665.

16. Costa,A.I., Koning,D., Ladell,K., Mclaren,J.E., Grady,B.P.X., Schellens,I.M.M., van Ham,P., Nijhuis,M., Borghans,J.A.M., Keşmir,C., et al. (2015) Complex T-Cell Receptor Repertoire Dynamics Underlie the CD8 + T-Cell Response to HIV-1. 89, 110–119.

17. Miconnet,I., Marrau,A., Farina,A., Taffé,P., Vigano,S., Harari,A. and Pantaleo,G. (2011) Large TCR Diversity of Virus-Specific CD8 T Cells Provides the Mechanistic Basis for Massive TCR Renewal after Antigen Exposure. J. Immunol., 186, 7039–7049.

18. Venturi,V., Chin,H.Y., Asher,T.E., Ladell,K., Scheinberg,P., Bornstein,E., van Bockel,D., Kelleher,A.D., Douek,D.C., Price,D.A., et al. (2008) TCR β-Chain Sharing in Human CD8+ T Cell Responses to Cytomegalovirus and EBV. J. Immunol., 181, 7853–7862.

19. Lefranc,M.P., Giudicelli,V., Duroux,P., Jabado-Michaloud,J., Folch,G., Aouinti,S., Carillon,E., Duvergey,H., Houles,A., Paysan-Lafosse,T., et al. (2015) IMGT, the international ImMunoGeneTics information system 25 years on. Nucleic Acids Res., 43, D413–D422.

20. Robins,H.S., Srivastava,S.K., Campregher,P. V, Turtle,C.J., Andriesen,J., Riddell,S.R., Carlson,C.S. and Warren,E.H. (2010) Overlap and effective size of the human CD8+ T-cell receptor repertoire. Sci. Transl. Med., 2, 47ra64.

21. Dean,J., Emerson,R.O., Vignali,M., Sherwood,A.M., Rieder,M.J., Carlson,C.S. and Robins,H.S. (2015) Annotation of pseudogenic gene segments by massively parallel sequencing of rearranged lymphocyte receptor loci. Genome Med., 7, 123.

22. Jokinen,E., Heinonen,M., Huuhtanen,J., Mustjoki,S. and Lähdesmäki,H. (2019) TCRGP□: Determining epitope specificity of T cell receptors. bioRxiv.

23. Pedregosa,F., Varoquaux,G., Gramfort,A., Michel,V., Thirion,B., Grisel,O., Blondel,M., Prettenhofer,P., Weiss,R., Dubourg,V., et al. (2011) Scikit-learn: Machine Learning in Python. J. Mach. Learn. Res., 12, 2825–2830.

24. Django Software Foundation (2018) Django (Version 2.0.7).

25. Jones,E., Oliphant,T., Peterson,P. and others (2001-) SciPy: Open source scientific tools for Python.

26. Goloborodko,A.A., Levitsky,L.I., Ivanov,M. V. and Gorshkov,M. V. (2013) Pyteomics — a Python Framework for Exploratory Data Analysis and Rapid Software Prototyping in Proteomics. J. Am. Soc. Mass Spectrom., 24, 301–304.

27. VanderPlas,J., Granger,B.E., Heer,J., Moritz,D., Wongsuphasawat,K., Satyanarayan,A., Lees,E., Timofeev,I., Welsh,B. and Sievert,S. (2018) Altair□: Interactive Statistical Visualizations for Python. J. Open Source Softw., 3, 1057.

28. Oliphant,T.E. (2006) A guide to NumPy. Trelgol Publishing USA.

29. McKinney,W. (2010) Data Structures for Statistical Computing in Python. In Proceedings of the 9th Python in Science Conference.pp. 51–56.

30. Dask Development Team (2016) Dask: Library for dynamic task scheduling.

31. Bolotin,D.A., Poslavsky,S., Mitrophanov,I., Shugay,M., Mamedov,I.Z., Putintseva,E. V. and Chudakov,D.M. (2015) MiXCR: Software for comprehensive adaptive immunity profiling. Nat. Methods, 12, 380–381.

32. Redmond,D., Poran,A. and Elemento,O. (2016) Single-cell TCRseq□: paired recovery of entire T-cell alpha and beta chain transcripts in T-cell receptors from single-cell RNAseq. Genome Med., 8, 80.

33. Lee,E.S., Thomas,P.G., Mold,J.E. and Yates,A.J. (2017) Identifying T Cell Receptors from High-Throughput Sequencing□: Dealing with Promiscuity in TCR α and TCR β Pairing. PLoS Comput. Biol., 13, e1005313.

