## supplemental materials for "TCRex: detection of enriched T cell epitope specificity in full T cell receptor sequence repertoires"

### Supplemental material S1: Standard TCRex filtering steps

The TCRex webtool performs a few filtering steps on each uploaded TCR dataset: (1) TCR beta sequences with orphon genes, (2) non-canonical TCR beta sequences (i.e. TCRs not starting with cysteine and/or not ending with phenylalanine) and (3) TCR beta sequences containing non-amino acid characters (e.g a stop codon) or lowercase amino acids are removed. As the immunoSEQ Analyzer format gives information on the frame type of each TCR beta sequence, TCRex retains only the TCR beta sequences from these files that are reported to be in-frame. The IMGT parser ensures that all V/J genes are presented in the IMGT format. This IMGT parser can be switched off when wanted. In this paper, we used the default option where the IMGT parser was activated. Duplicate TCR beta sequences are automatically removed by TCRex. Therefore, all TCR repertoire sizes used in the calculations of the percentage of epitope-specific T cells and the enrichment analyses in this paper refer to the number of unique epitope-specific TCR beta sequences that passed all filtering steps.

### Supplemental material S2: web tool instructions

To start a prediction analysis, users are required to upload a file containing TCR data. Different file formats are supported by TCRex, including the immunoSEQ Analyzer (https://www.adaptivebiotech.com/immunoseq) and MiXCR (1) output formats. In addition, we propose a simple tab-delimited format that includes the CDR3 amino acid sequence and the V/J genes for all TCR beta sequences following the international ImMunoGeneTics information system (IMGT) notation (2). In case no V/J gene information is available, users are advised to provide the corresponding V/J families.

After uploading the input TCR data file, one or more epitopes can be selected from the database. In this release (version 0.3.0), prediction models for 49 different epitopes are available, including 44 viral and 5 cancer epitopes. With the development and improvement of new TCR sequencing techniques and the rising interest in epitope-specific TCR repertoire analysis, we expect this number to grow rapidly in the near future. The database will be updated regularly with new epitope data made available in the scientific literature. Alternatively, it is also possible to make predictions for epitopes that are not available in the database. To this end, users can upload both their own dataset containing epitope-specific TCR sequences for a single epitope and a TCR test dataset. Both files must follow one of the data formats described above. The epitope-specific TCR sequences are then used to train a new prediction model and the resulting model is subsequently used to make predictions on the uploaded test dataset. To ensure data privacy, these custom models are not made available to other users.

Thereafter, the user can choose whether the V/J genes in the input format need to be changed to the standard IMGT format and select a threshold for the enrichment analysis. This threshold represents the percentage of epitope-specific TCRs in a background dataset. By default, a value of 0.01% is chosen.

When all the required information is submitted, the web tool redirects the user to a web page that gives an overview of all the steps in the prediction process and the current status of the analysis. Once the analysis is finalized, an interactive results summary of the prediction results is given. This allows the user to select a BPR threshold to filter the prediction results. By default, a threshold of 0.01% BPR is used. The filtered results are made available for download as a tab-delimited file. For every TCR sequence provided by the user and for every selected epitope, this file contains a score that represents the probability of epitope-TCR recognition. In addition, each epitope-TCR pair is supplemented with a BPR value, which is used to filter the results. In case the user trained a new prediction model, the results are supplemented with a summary of the performance metrics along with the ROC and PR curves and a visualization of the important features of the learned model. All results are kept available for seven days.

### Supplemental material S3: Performance for all trained epitope-specific prediction models

**Table S1: Performance metrics of the trained prediction models.** For each model, the size of the positive training dataset, the balanced accuracy, the area under the ROC curve (AUC) and the average precision are given. The last column indicates whether the trained model passed all performance criteria.

| Source | Epitope | Size positive training dataset | Balanced accuracy | AUC | Average precision | Passed performance criteria |
|---|---|---|---|---|---|---|
| CMV | IPSINVHHY | 85 | 0.7 ± 0.02 | 0.84 ± 0.04 | 0.66 ± 0.06 | Yes |
| CMV | MLNIPSINV | 73 | 0.51 ± 0.01 | 0.61 ± 0.04 | 0.25 ± 0.04 | No |
| CMV | NLVPMVATV | 4812 | 0.56 ± 0.0 | 0.72 ± 0.01 | 0.39 ± 0.01 | Yes |
| CMV | QIKVRVKMV | 36 | 0.5 ± 0.0 | 0.79 ± 0.05 | 0.4 ± 0.09 | Yes |
| CMV | QYDPVAALF | 41 | 0.7 ± 0.06 | 0.82 ± 0.06 | 0.6 ± 0.06 | Yes |
| CMV | TPRVTGGGAM | 258 | 0.7 ± 0.02 | 0.88 ± 0.03 | 0.75 ± 0.04 | Yes |
| CMV | VTEHDTLLY | 277 | 0.54 ± 0.0 | 0.78 ± 0.02 | 0.39 ± 0.04 | Yes |
| CMV | YSEHPTFTSQY | 74 | 0.59 ± 0.06 | 0.93 ± 0.04 | 0.71 ± 0.06 | Yes |
| DENV1 | GTSGSPIVNR | 165 | 0.75 ± 0.05 | 0.88 ± 0.04 | 0.76 ± 0.06 | Yes |
| DENV2 | GTSGSPIIDK | 60 | 0.61 ± 0.06 | 0.74 ± 0.08 | 0.49 ± 0.04 | Yes |
| DENV3/4 | GTSGSPIINR | 158 | 0.73 ± 0.02 | 0.86 ± 0.03 | 0.75 ± 0.03 | Yes |
| EBV | EPLPQGQLTAY | 36 | 0.64 ± 0.08 | 0.91 ± 0.06 | 0.68 ± 0.18 | Yes |
| EBV | GLCTLVAML | 1208 | 0.66 ± 0.01 | 0.82 ± 0.02 | 0.59 ± 0.0 | Yes |
| EBV | HPVGEADYFEY | 32 | 0.72 ± 0.08 | 0.86 ± 0.08 | 0.68 ± 0.13 | Yes |
| EBV | IVTDFSVIK | 46 | 0.6 ± 0.05 | 0.83 ± 0.04 | 0.53 ± 0.05 | Yes |
| EBV | RAKFKQLL | 262 | 0.65 ± 0.02 | 0.89 ± 0.01 | 0.69 ± 0.04 | Yes |
| EBV | YVLDHLIVV | 103 | 0.52 ± 0.02 | 0.76 ± 0.07 | 0.44 ± 0.09 | Yes |
| HCV | ARMILMTHF | 66 | 0.74 ± 0.07 | 0.86 ± 0.08 | 0.69 ± 0.12 | Yes |
| HCV | ATDALMTGY | 177 | 0.75 ± 0.03 | 0.91 ± 0.04 | 0.77 ± 0.06 | Yes |
| HCV | CINGVCWTV | 131 | 0.53 ± 0.03 | 0.75 ± 0.06 | 0.37 ± 0.11 | Yes |
| HCV | HSKKKCDEL | 45 | 0.77 ± 0.09 | 0.99 ± 0.01 | 0.93 ± 0.03 | Yes |
| HCV | KLVALGINAV | 65 | 0.62 ± 0.05 | 0.73 ± 0.08 | 0.51 ± 0.14 | Yes |
| HIV | EIYKRWII | 185 | 0.65 ± 0.02 | 0.75 ± 0.06 | 0.52 ± 0.04 | Yes |
| HIV | FLKEKGGL | 156 | 0.6 ± 0.03 | 0.76 ± 0.06 | 0.48 ± 0.1 | Yes |
| HIV | FPRPWLHGL | 120 | 0.81 ± 0.02 | 0.93 ± 0.02 | 0.83 ± 0.03 | Yes |
| HIV | FRDYVDRFYKTLRAEQASQE | 367 | 0.87 ± 0.02 | 1.0 ± 0.0 | 0.96 ± 0.01 | Yes |
| HIV | GPGHKARVL | 66 | 0.51 ± 0.02 | 0.67 ± 0.06 | 0.31 ± 0.11 | No |
| HIV | HPKVSSEVHI | 75 | 0.69 ± 0.07 | 0.87 ± 0.08 | 0.7 ± 0.16 | Yes |
| HIV | IIKDYGKQM | 54 | 0.81 ± 0.07 | 0.95 ± 0.04 | 0.84 ± 0.07 | Yes |
| HIV | ISPRTLNAW | 58 | 0.61 ± 0.06 | 0.84 ± 0.03 | 0.6 ± 0.12 | Yes |
| HIV | KAFSPEVIPMF | 210 | 0.73 ± 0.02 | 0.9 ± 0.01 | 0.71 ± 0.03 | Yes |
| HIV | KRWIILGLNK | 396 | 0.65 ± 0.02 | 0.85 ± 0.04 | 0.64 ± 0.08 | Yes |
| HIV | KRWIIMGLNK | 75 | 0.69 ± 0.05 | 0.85 ± 0.09 | 0.73 ± 0.12 | Yes |
| HIV | LPPIVAKEI | 62 | 0.79 ± 0.04 | 0.9 ± 0.05 | 0.78 ± 0.06 | Yes |
| HIV | QASQEVKNW | 31 | 0.58 ± 0.05 | 0.6 ± 0.11 | 0.31 ± 0.12 | No |
| HIV | QVPLRPMTYK | 48 | 0.64 ± 0.04 | 0.77 ± 0.11 | 0.51 ± 0.16 | Yes |
| HIV | RFYKTLRAEQASQ | 210 | 0.9 ± 0.03 | 0.99 ± 0.0 | 0.96 ± 0.01 | Yes |
| HIV | RLRPGGKKK | 31 | 0.73 ± 0.06 | 0.89 ± 0.09 | 0.73 ± 0.2 | Yes |
| HIV | RYPLTFGWCF | 30 | 0.52 ± 0.03 | 0.72 ± 0.13 | 0.42 ± 0.12 | Yes |
| HIV | SLYNTVATL | 58 | 0.63 ± 0.03 | 0.59 ± 0.06 | 0.4 ± 0.05 | No |
| HIV | TPGPGVRYPL | 86 | 0.68 ± 0.04 | 0.88 ± 0.03 | 0.74 ± 0.07 | Yes |
| HIV | TPQDLNTML | 159 | 0.81 ± 0.02 | 0.94 ± 0.04 | 0.87 ± 0.06 | Yes |
| HSV2 | RPRGEVRFL | 63 | 0.77 ± 0.04 | 0.92 ± 0.05 | 0.84 ± 0.08 | Yes |
| HTLV1 | SFHSLHLLF | 131 | 0.68 ± 0.03 | 0.81 ± 0.08 | 0.63 ± 0.08 | Yes |
| Cancer | NLSALGIFST | 111 | 0.51 ± 0.01 | 0.66 ± 0.04 | 0.19 ± 0.03 | No |
| Influenza | GILGFVFTL | 3107 | 0.68 ± 0.01 | 0.81 ± 0.01 | 0.59 ± 0.03 | Yes |
| Influenza | LPRRSGAAGA | 2137 | 0.54 ± 0.01 | 0.84 ± 0.01 | 0.4 ± 0.02 | Yes |
| Influenza | PKYVKQNTLKLAT | 292 | 0.52 ± 0.01 | 0.72 ± 0.02 | 0.36 ± 0.05 | Yes |
| Melanoma | AMFWSVPTV | 82 | 0.59 ± 0.04 | 0.78 ± 0.04 | 0.48 ± 0.12 | Yes |
| Melanoma | EAAGIGILTV | 266 | 0.69 ± 0.03 | 0.9 ± 0.01 | 0.74 ± 0.01 | Yes |

| | | | | | | |
|---|---|---|---|---|---|---|
| Melanoma | ELAGIGILTV | 1035 | 0.54 ± 0.0 | 0.72 ± 0.01 | 0.37 ± 0.03 | Yes |
| Melanoma | FLYNLLTRV | 61 | 0.63 ± 0.08 | 0.87 ± 0.03 | 0.59 ± 0.1 | Yes |
| Multiple Myeloma | LLLGIGILV | 233 | 0.58 ± 0.01 | 0.77 ± 0.04 | 0.44 ± 0.06 | Yes |
| YFV | LLWNGPMAV | 474 | 0.66 ± 0.01 | 0.79 ± 0.01 | 0.53 ± 0.03 | Yes |

### Supplemental material S4: MHC background of epitope-specific models

Table S2 gives an overview of the MHC background of all epitopes listed in table S1. For each epitope, it reports the MHC molecules that were associated with the collected TCR beta sequences. Since some of the epitope-specific TCR sequences were collected multiple times due to overlap within or between different sources, we checked whether all these TCRs had the same MHC background. In case multiple MHC molecules were associated with the same TCR, all MHC molecules are reported in the same row and separated with a comma. The third column represents the number of unique epitope-specific TCRs that correspond with the MHC background in the second column. The last column gives an overview of the total number of unique TCRs for each epitope, which is identical to the positive training size shown in table S1.

**Table S2: Overview of the MHC background of all epitope-specific TCRs in the training dataset**

| Epitope | MHC background | Number of unique TCR sequences | Total training size per epitope |
|---|---|---|---|
| AMFWSVPTV | HLA-A*02:01 | 82 | 82 |
| ARMILMTHF | HLA-B*27 | 66 | 66 |
| ATDALMTGY | HLA-A*01 | 34 | 177 |
| | HLA-A*01:01 | 143 | |
| CINGVCWTV | HLA-A*02 | 58 | 131 |
| | HLA-A*02, HLA-A*02:01 | 2 | |
| | HLA-A*02:01 | 71 | |
| EAAGIGILTV | HLA-A*02 | 49 | 266 |
| | HLA-A*02:01 | 210 | |
| | HLA-A*02:01:48 | 1 | |
| | unknown | 6 | |
| EIYKRWII | HLA-B*08 | 63 | 185 |
| | HLA-B*08, HLA-B*08:01 | 113 | |
| | HLA-B*08:01 | 9 | |
| ELAGIGILTV | HLA-A*02 | 968 | 1035 |
| | HLA-A*02, HLA-A*02:01, HLA-A2 | 1 | |
| | HLA-A*02, HLA-A2 | 4 | |
| | HLA-A*02:01 | 45 | |
| | HLA-A*02:01:48 | 4 | |
| | HLA-A2 | 13 | |
| EPLPQGQLTAY | HLA-B*35:01 | 33 | 36 |
| | HLA-B*35:01, HLA-B*35:02 | 1 | |
| | HLA-B*35:01, HLA-B*35:42:01 | 1 | |
| | HLA-B*35:02 | 1 | |
| FLKEKGGL | HLA-B*08 | 73 | 156 |
| | HLA-B*08, HLA-B*08:01 | 67 | |

| | | | |
|---|---|---|---|
| | HLA-B*08:01 | 16 | |
| FLYNLLTRV | HLA-A*02:01 | 61 | 61 |
| FPRPWLHGL | HLA-B*42:01 | 120 | 120 |
| FRDYVDRFYKTLRAEQASQE | HLA-DRA*01&HLA-DRB1*01:01 | 3 | 367 |
| | HLA-DRA*01&HLA-DRB1*01:01, HLA-DRA*01&HLA-DRB1*07:01 | 5 | |
| | HLA-DRA*01&HLA-DRB1*01:01, HLA-DRA*01&HLA-DRB1*07:01, HLA-DRA*01&HLA-DRB1*15:02 | 1 | |
| | HLA-DRA*01&HLA-DRB1*07:01 | 12 | |
| | HLA-DRA*01&HLA-DRB1*11:01 | 4 | |
| | HLA-DRA*01&HLA-DRB1*11:01, HLA-DRA*01&HLA-DRB1*15:02, HLA-DRA*01&HLA-DRB5*01:01, HLA-DRA*01:01&HLA-DRB1*11:01, HLA-DRA*01:01&HLA-DRB1*15:01, HLA-DRA*01:01&HLA-DRB1*15:02, HLA-DRA*01:01&HLA-DRB5*01:01 | 1 | |
| | HLA-DRA*01&HLA-DRB1*11:01, HLA-DRA*01&HLA-DRB5*01:01 | 1 | |
| | HLA-DRA*01&HLA-DRB1*11:01, HLA-DRA*01&HLA-DRB5*01:01, HLA-DRA*01:01&HLA-DRB1*11:01, HLA-DRA*01:01&HLA-DRB5*01:01 | 1 | |
| | HLA-DRA*01&HLA-DRB1*15:02 | 1 | |
| | HLA-DRA*01&HLA-DRB1*15:02, HLA-DRA*01&HLA-DRB5*01:01 | 2 | |
| | HLA-DRA*01&HLA-DRB1*15:02, HLA-DRA*01&HLA-DRB5*01:01, HLA-DRA*01:01&HLA-DRB5*01:01 | 1 | |
| | HLA-DRA*01&HLA-DRB5*01:01 | 29 | |
| | HLA-DRA*01:01&HLA-DRB1*01:01 | 29 | |
| | HLA-DRA*01:01&HLA-DRB1*01:01, HLA-DRA*01:01&HLA-DRB1*15:01 | 3 | |
| | HLA-DRA*01:01&HLA-DRB1*01:01, HLA-DRA*01:01&HLA-DRB5*01:01 | 2 | |
| | HLA-DRA*01:01&HLA-DRB1*11:01 | 68 | |
| | HLA-DRA*01:01&HLA-DRB1*11:01, HLA-DRA*01:01&HLA-DRB1*15:01, HLA-DRA*01:01&HLA-DRB5*01:01 | 1 | |
| | HLA-DRA*01:01&HLA-DRB1*11:01, HLA-DRA*01:01&HLA-DRB5*01:01 | 9 | |
| | HLA-DRA*01:01&HLA-DRB1*15:01 | 51 | |
| | HLA-DRA*01:01&HLA-DRB1*15:01, HLA-DRA*01:01&HLA-DRB5*01:01 | 13 | |
| | HLA-DRA*01:01&HLA-DRB1*15:02, HLA-DRA*01:01&HLA-DRB5*01:01 | 2 | |
| | HLA-DRA*01:01&HLA-DRB5*01:01 | 128 | |
| GILGFVFTL | HLA-A*02 | 1805 | 3107 |
| | HLA-A*02, HLA-A*02:01 | 59 | |
| | HLA-A*02, HLA-A*02:01, HLA-A*02:01:48 | 1 | |
| | HLA-A*02, HLA-A*02:01, HLA-A*02:01:48, HLA-A2 | 1 | |
| | HLA-A*02, HLA-A*02:01, HLA-A2 | 41 | |
| | HLA-A*02, HLA-A2 | 204 | |
| | HLA-A*02:01 | 568 | |

| | | | |
|---|---|---|---|
| | HLA-A*02:01, HLA-A2 | 268 | |
| | HLA-A2 | 160 | |
| GLCTLVAML | HLA-A*02 | 204 | 1208 |
| | HLA-A*02, HLA-A*02:01 | 16 | |
| | HLA-A*02, HLA-A*02:01, HLA-A*02:01:48, HLA-A2 | 1 | |
| | HLA-A*02, HLA-A*02:01, HLA-A*2:01, HLA-A2 | 1 | |
| | HLA-A*02, HLA-A*02:01, HLA-A2 | 12 | |
| | HLA-A*02:01 | 332 | |
| | HLA-A*02:01, HLA-A2 | 439 | |
| | HLA-A*2:01 | 6 | |
| | HLA-A2 | 197 | |
| GPGHKARVL | HLA-B*07:02 | 66 | 66 |
| GTSGSPIIDK | HLA-A*11:01 | 60 | 60 |
| GTSGSPIINR | HLA-A*11:01 | 158 | 158 |
| GTSGSPIVNR | HLA-A*11:01 | 165 | 165 |
| HPKVSSEVHI | HLA-B*42:01 | 75 | 75 |
| HPVGEADYFEY | HLA-B*35:01 | 23 | 32 |
| | HLA-B*35:01, HLA-B*35:08, HLA-B*35:08:01, HLA-B*35:42:01 | 1 | |
| | HLA-B*35:08 | 8 | |
| HSKKKCDEL | HLA-B*08:01 | 44 | 45 |
| | HLA-B*08:01, HLA-B*08:01:29 | 1 | |
| IIKDYGKQM | HLA-B*42:01 | 54 | 54 |
| IPSINVHHY | HLA-B*35 | 10 | 85 |
| | HLA-B*35:01 | 75 | |
| ISPRTLNAW | HLA-B*57 | 4 | 58 |
| | HLA-B*57, HLA-B*57:01, HLA-B*57:03 | 1 | |
| | HLA-B*57, HLA-B*57:03 | 5 | |
| | HLA-B*57:01 | 44 | |
| | HLA-B*57:03 | 4 | |
| IVTDFSVIK | HLA-A*011 | 6 | 46 |
| | HLA-A*011, HLA-A*11 | 3 | |
| | HLA-A*11 | 6 | |
| | HLA-A*11:01 | 31 | |
| KAFSPEVIPMF | HLA-B*57 | 9 | 210 |
| | HLA-B*57, HLA-B*57:01 | 33 | |
| | HLA-B*57, HLA-B*57:01, HLA-B*57:03 | 4 | |
| | HLA-B*57:01 | 133 | |
| | HLA-B*57:01, HLA-B*57:03 | 16 | |
| | HLA-B*57:01, HLA-B*57:06 | 1 | |
| | HLA-B*57:03 | 14 | |
| KLVALGINAV | HLA-A*02 | 65 | 65 |

| | | | |
|---|---|---|---|
| KRWIILGLNK | HLA-B*27 | 66 | 396 |
| | HLA-B*27, HLA-B*27:05 | 54 | |
| | HLA-B*27:05 | 275 | |
| | HLA-B*27:05:31 | 1 | |
| KRWIIMGLNK | HLA-B*27 | 7 | 75 |
| | HLA-B*27:05 | 67 | |
| | HLA-B*27:05:31 | 1 | |
| LLLGIGILV | HLA-A*02 | 233 | 233 |
| LLWNGPMAV | HLA-A*02 | 138 | 474 |
| | HLA-A*02, HLA-A*02:01 | 1 | |
| | HLA-A*02:01 | 335 | |
| LPPIVAKEI | HLA-B*42:01 | 62 | 62 |
| LPRRSGAAGA | HLA-B*07:02 | 1 | 2137 |
| | HLA-B*07:02, HLA-B7 | 158 | |
| | HLA-B7 | 1978 | |
| MLNIPSINV | HLA-A*02 | 73 | 73 |
| NLSALGIFST | HLA-A*02 | 111 | 111 |
| NLVPMVATV | HLA-A*02 | 4066 | 4812 |
| | HLA-A*02, HLA-A*02:01 | 19 | |
| | HLA-A*02, HLA-A*02:01, HLA-A*02:01:59, HLA-A2 | 1 | |
| | HLA-A*02, HLA-A*02:01, HLA-A2 | 2 | |
| | HLA-A*02, HLA-A2 | 3 | |
| | HLA-A*02:01 | 581 | |
| | HLA-A*02:01, HLA-A2 | 7 | |
| | HLA-A*02:01:110 | 1 | |
| | HLA-A*02:01:98 | 1 | |
| | HLA-A2 | 126 | |
| | unknown | 5 | |
| PKYVKQNTLKLAT | HLA-DRA*01&HLA-DRB1*01 | 107 | 292 |
| | HLA-DRA*01:01&HLA-DRB1*04:01 | 183 | |
| | HLA-DRA*01:02:03&HLA-DRB1*01:01:01 | 1 | |
| | HLA-DRA*01:02:03&HLA-DRB1*01:01:01, HLA-DRA*01:02:03&HLA-DRB1*04:01:01 | 1 | |
| QASQEVKNW | HLA-B*53 | 8 | 31 |
| | HLA-B*53, HLA-B*58 | 1 | |
| | HLA-B*57 | 5 | |
| | HLA-B*57:01 | 16 | |
| | HLA-B*58 | 1 | |
| QIKVRVKMV | HLA-B*08:01 | 36 | 36 |
| QVPLRPMTYK | HLA-A*03 | 28 | 48 |
| | HLA-A*03, HLA-A*11 | 5 | |
| | HLA-A*03:01 | 10 | |

| | | | |
|---|---|---|---|
| | HLA-A*11 | 5 | |
| QYDPVAALF | HLA-A*24:02 | 41 | 41 |
| RAKFKQLL | HLA-B*08:01 | 260 | 262 |
| | HLA-B*8 | 2 | |
| RFYKTLRAEQASQ | HLA-DR1 | 16 | 210 |
| | HLA-DR1, HLA-DR15 | 3 | |
| | HLA-DR1, HLA-DR5 | 2 | |
| | HLA-DR11 | 47 | |
| | HLA-DR11, HLA-DR15, HLA-DR5 | 1 | |
| | HLA-DR11, HLA-DR5 | 3 | |
| | HLA-DR15 | 100 | |
| | HLA-DR15, HLA-DR5 | 6 | |
| | HLA-DR5 | 32 | |
| RLRPGGKKK | HLA-A*03:01 | 31 | 31 |
| RPRGEVRFL | HLA-B*07:02 | 63 | 63 |
| RYPLTFGWCF | HLA-A*24:02 | 29 | 30 |
| | HLA-A*24:02:84 | 1 | |
| SFHSLHLLF | HLA-A*24:02 | 131 | 131 |
| SLYNTVATL | HLA-A*02 | 1 | 58 |
| | HLA-A*02, HLA-A*02:01 | 21 | |
| | HLA-A*02:01 | 36 | |
| TPGPGVRYPL | HLA-B*07:02 | 23 | 86 |
| | HLA-B*42:01 | 63 | |
| TPQDLNTML | HLA-B*42 | 5 | 159 |
| | HLA-B*42:01 | 114 | |
| | HLA-B*81:01 | 40 | |
| TPRVTGGGAM | HLA-B*07 | 18 | 258 |
| | HLA-B*07, HLA-B*07:02, HLA-B7 | 1 | |
| | HLA-B*07:02 | 208 | |
| | HLA-B*07:02, HLA-B7 | 19 | |
| | HLA-B7 | 12 | |
| VTEHDTLLY | HLA-A*01:01, HLA-A1 | 201 | 277 |
| | HLA-A1 | 76 | |
| YSEHPTFTSQY | HLA-A*01:01 | 74 | 74 |
| YVLDHLIVV | HLA-A*02 | 10 | 103 |
| | HLA-A*02:01 | 93 | |

### Supplemental material S5: data selection for the leave-on-study-out validation

A leave-one-study-out validation strategy was carried out to assess whether our epitope-specific models are suitable to perform predictions for new studies. This analysis was restricted to viral epitopes for which a large amount of training data (at least 1000 different TCR beta sequences) was collected from different sources. Only three epitopes fulfilled these criteria: NLVPMVATV, GILGFVFTL and GLCTLVAML. For each of these epitopes, we trained epitope-specific models using all collected data except for the TCR beta sequences from one study. The latter was used as an external validation dataset to evaluate the performance of the model on new data. This left out study was selected based on the reported number of epitope-specific TCR cells: (1) studies which reported a limited number of TCR sequences were not eligible as they do not give a good idea of the identification rate of a normal study (2) studies with a large amount of epitope-specific TCRs were also not eligible since we wanted to have a large training dataset. Tables S3-S5 give an overview of the number of epitope-specific TCRs collected from each study following the quality filtering steps as explained in the main text. These numbers are an estimate of the true data size of the studies as duplicate TCRs might still be present, due to an overlap of the different databases and sources we used, and the data were not subjected to the parsing steps that are automatically performed by the TCRex webtool. The selected studies are highlighted in green. All have estimated sizes of approximately 150 TCR sequences.

**Table S3: Overview of the references containing GLCTLVAML-specific TCRs**

| Reference | Number of TCR beta sequences |
|---|---|
| PMID:28636589 | 1103 |
| PMID:28636592 | 132 |
| https://github.com/antigenomics/vdjdb-db/issues/243 | 130 |
| PMID:19017975 | 92 |
| PMID:12504586 | 89 |
| PMID:21555537 | 44 |
| PMID:10925283 | 39 |
| PMID:24512815 | 35 |
| PMID:11046006 | 33 |
| PMID:19542443 | 23 |
| PMID:21124993 | 19 |
| PMID:23267020 | 17 |
| PMID:25339770 | 12 |
| PMID:27645996 | 11 |
| PMID:11592365 | 7 |
| PMID:16287711 | 4 |
| PMID:25801351 | 4 |

**Table S4: Overview of the references containing NLVPMVATV-specific TCRs**

| Reference | Number of TCR beta sequences |
|---|---|
| PMID:28423320 | 4049 |
| PMID:19017975 | 172 |
| PMID:28636592 | 152 |
| PMID:28636589 | 108 |
| PMID:21555537 | 87 |
| PMID:21374820 | 63 |
| PMID:16287711 | 57 |
| PMID:16237109 | 43 |
| PMID:28623251 | 33 |
| https://github.com/antigenomics/vdjdb-db/issues/252 | 29 |
| PMID:24711416 | 28 |
| PMID:19014475 | 25 |
| PMID:25801351 | 20 |
| PMID:25925682 | 13 |
| PMID:11756174 | 9 |
| PMID:9971792 | 9 |
| PMID:19542443 | 9 |
| PMID:24512815 | 7 |
| PMID:24069285 | 7 |
| PMID:23267020 | 6 |
| PMID:17709536 | 5 |
| PMID:19403059 | 5 |
| PMID:25576336 | 5 |
| PMID:25339770 | 4 |
| PMID:12616496 | 3 |
| PMID:21135165 | 3 |
| PMID:26429912 | 2 |
| PMID:11930311 | 2 |
| PMID:19542454 | 1 |
| PMID:19864595 | 1 |

**Table S5: Overview of the references containing GILGFVFTL-specific TCRs**

| Reference | Number of TCR beta sequences |
|---|---|
| PMID:28423320 | 2295 |
| PMID:28636589 | 775 |
| PMID:28636592 | 406 |
| PMID:28300170 | 153 |
| PMID:25609818 | 101 |
| PMID:29483513 | 32 |
| PMID:27645996 | 29 |
| PMID:25339770 | 26 |
| PMID:28250417 | 22 |
| PMID:25801351 | 16 |
| PMID:7807026 | 12 |
| PMID:18275829 | 1 |
| PMID:12796775 | 1 |

**Supplemental material S6: Calculation of the performance metrics for the leave-on-study-out validation**

Following standard formulas were used to calculate the performance metrics in table 1 of the main text.

$$Accuracy = \frac{TP + TN}{TP + TN + FP + FN}$$

$$Sensitivity = \frac{TP}{TP + FN}$$

$$Specificity = \frac{TN}{TN + FP}$$

$$Precision = \frac{TP}{TP + FP}$$

With:

- FP = number of identified TCRs in the cancer dataset
- TN = number of TCRs in the cancer dataset that were classified as non-epitope binding
- TP = number of identified TCRs in the epitope-specific external validation dataset
- FN = number of TCRs in the epitope-specific external validation dataset that were classified as non-epitope binding

### Supplemental material S7: Public versus TCRex identified LLWNGPMAV-specific TCRs

**Table S6: Number of LLWNGPMAV-specific TCR sequences in the post-vaccination PBMC samples of (3).** For each volunteer from (3) are given: the number of LLWNGPMAV-specific TCRs in the post-vaccination repertoire identified with TCRex (0.01% BPR), the number of training LLWNGPMAV-specific TCRs in the post-vaccination repertoire and the corresponding overlap (i.e. the number of training TCR sequences that were identified with TCRex).

| Volunteer | TCRs identified by TCRex | Training TCRs | Overlap |
|---|---|---|---|
| 1 | 32 | 13 | 4 |
| 2 | 65 | 19 | 7 |
| 3 | 87 | 34 | 9 |
| 4 | 34 | 11 | 2 |
| 5 | 63 | 20 | 7 |
| 6 | 32 | 12 | 4 |
| 7 | 11 | 7 | 2 |
| 8 | 27 | 12 | 4 |
| 9 | 18 | 6 | 1 |

**Table S7: Number of LLWNGPMAV-specific TCR sequences in the post-vaccination PBMC samples of (4).** For each volunteer from (4) are given: the number of LLWNGPMAV-specific TCRs in the post-vaccination repertoire identified with TCRex (0.01% BPR), the number of training LLWNGPMAV-specific TCRs in the post-vaccination repertoire and the corresponding overlap (i.e. the number of training TCR sequences that were identified with TCRex).

| Volunteer | TCRs identified by TCRex | Training TCRs | Overlap |
|---|---|---|---|
| P1 | 92 | 31 | 12 |
| P2 | 110 | 36 | 9 |
| Q1 | 79 | 26 | 12 |
| Q2 | 92 | 33 | 12 |
| S1 | 170 | 48 | 21 |
| S2 | 273 | 53 | 26 |

### References

1. Bolotin,D.A., Poslavsky,S., Mitrophanov,I., Shugay,M., Mamedov,I.Z., Putintseva,E. V. and Chudakov,D.M. (2015) MiXCR: Software for comprehensive adaptive immunity profiling. *Nat. Methods*, **12**, 380–381.
2. Lefranc,M.P., Giudicelli,V., Duroux,P., Jabado-Michaloud,J., Folch,G., Aouinti,S., Carillon,E., Duvergey,H., Houles,A., Paysan-Lafosse,T., *et al.* (2015) IMGT, the international ImMunoGeneTics information system 25 years on. *Nucleic Acids Res.*, **43**, D413–D422.
3. Dewitt,W.S., Emerson,R.O., Lindau,P., Vignali,M., Snyder,T.M., Desmarais,C., Sanders,C., Utsugi,H., Warren,E.H., McElrath,J., *et al.* (2015) Dynamics of the Cytotoxic T Cell Response to a Model of Acute Viral Infection. *J. Virol.*, **89**, 4517–4526.
4. Pogorelyy,M. V., Minervina,A.A., Touzel,M.P., Sycheva,A.L., Komech,E.A., Kovalenko,E.I., Karganova,G.G., Egorov,E.S., Komkov,A.Y., Chudakov,D.M., *et al.* (2018) Precise tracking of vaccine-responding T cell clones reveals convergent and personalized response in identical twins. *PNAS*, **115**, 12704–12709.
